# Antisense oligonucleotides targeting *UBE3A-ATS* restore expression of *UBE3A* by relieving transcriptional interference

**DOI:** 10.1101/2021.07.09.451826

**Authors:** Noelle D. Germain, Dea Gorka, Ryan Drennan, Paymaan Jafar-nejad, Amanda Whipple, Leighton Core, Eric S. Levine, Frank Rigo, Stormy J. Chamberlain

## Abstract

Angelman syndrome (AS) is a rare neurodevelopmental disorder caused by loss of function of the maternally inherited *UBE3A* allele. In neurons, the paternal allele of *UBE3A* is silenced in *cis* by the long noncoding RNA, *UBE3A-ATS*. Unsilencing paternal *UBE3A* by reducing *UBE3A-ATS* is a promising therapeutic approach for the treatment of AS. Here we show that targeted cleavage of *UBE3A-ATS* using antisense oligonucleotides (ASOs) restores *UBE3A* and rescues electrophysiological phenotypes in human AS neurons. We demonstrate that cleavage of *UBE3A-ATS* results in termination of its transcription by displacement of RNA Polymerase II. Reduced transcription of *UBE3A-ATS* allows transcription of *UBE3A* to proceed to completion, providing definitive evidence for the transcriptional interference model of paternal *UBE3A* silencing. These insights into the mechanism by which ASOs restore *UBE3A* inform the future development of nucleotide-based approaches for the treatment of AS, including alternative strategies for cleaving *UBE3A-ATS* that can be developed for long-term restoration of UBE3A function.

## Introduction

Angelman syndrome (AS), a rare neurodevelopmental disorder affecting ~ 1 in 15,000 people, is characterized by severe intellectual disability, absent speech, seizures, ataxia, and a happy demeanor ^1,2^. AS is caused by loss-of-function of the maternally inherited allele of *UBE3A,* which encodes an E3 ubiquitin ligase that targets proteins for degradation by the proteasome^3,4^. AS occurs as a result of large deletions of the q11-q13 region on maternally inherited chromosome 15, paternal uniparental disomy (UPD) of chromosome 15, imprinting defects, or deleterious mutations in the maternal *UBE3A* allele^3^. In non-neuronal tissues, *UBE3A* is expressed from both parental alleles. However, in neurons, a long non-coding antisense transcript (*UBE3A-ATS*), originating from the *SNURF-SNRPN* promoters, silences paternally inherited *UBE3A*^5,6^. Therefore, loss-of-function from the maternal allele leads to nearly complete loss of *UBE3A* RNA and protein in neurons. Activation of the intact, but transcriptionally silenced, paternal allele of *UBE3A* is an attractive and promising therapeutic target for AS since it could potentially restore *UBE3A* expression and function.

Studies in the AS mouse model have shed light on the mechanism whereby *Ube3a-ats* silences paternal *Ube3a*^7,8^. These studies determined that transcription of *Ube3a-ats* in *cis* is necessary for *Ube3a* imprinting. This “transcriptional interference model” purports that active transcription of *UBE3A-ATS* on the plus strand prevents full transcription of *UBE3A* from the minus strand by obstructing RNA Polymerase II (RNA Pol II). Work from our lab provided further support for the transcriptional interference model and demonstrated that by drastically increasing *UBE3A-ATS* transcription in human AS induced pluripotent stem cells (iPSCs), we could completely silence paternal *UBE3A* in non-neuronal cells^9^. Furthermore, the Beaudet lab inserted a transcriptional stop cassette within murine *Ube3a-ats* and found that it unsilenced *Ube3a* in the AS mouse model and rescued some of the AS behavioral phenotypes^8^. Subsequently, topoisomerase inhibitors and antisense oligonucleotides (ASOs) were both shown to reduce *Ube3a-ats* and restore *Ube3a* in mouse, providing proof-of-concept for reduction of *UBE3A-ATS* as a therapeutic approach to treat AS^10,11^.

ASOs are single-stranded oligonucleotide sequences that can bind to and target specific RNAs for degradation. ASOs reduce gene products by sterically inhibiting transcription or translation, by triggering RNase H-mediated degradation of the target RNA, or by inhibiting RNA splicing^12,13^. Studies of *Ube3a-ats* targeting ASOs in AS mouse neurons showed that only ASOs triggering an RNase H response were effective at restoring paternal *Ube3a* expression^11^. The 5’-> 3’ exonuclease XRN2 is required for the depletion of RNase H cleavage products generated after ASO binding to pre-mRNAs and nuclear retained RNAs^14,15^. XRN2 is implicated in the torpedo model of transcription termination where it degrades remaining RNA cleavage products and disengages RNA Pol II to terminate transcription^16^. Two elegant studies recently confirmed that ASOs that induce RNase H-mediated cleavage of target RNAs can cause transcription termination downstream of the cleavage site via displacement of RNA Pol II ^17,18^. However, this mechanism has not yet been confirmed for ASOs targeting *UBE3A-ATS*. Since ASOs targeting *UBE3A-ATS* are currently under early stages of investigation in clinical trials for the treatment of AS, it is important to provide these mechanistic data to guide development of optimal ASOs, as well as additional approaches to unsilence paternal *UBE3A* in AS.

Here, we show that ASOs targeting human *UBE3A-ATS* effectively rescue *UBE3A* expression, UBE3A protein levels, and electrophysiological deficits in our established human iPSC-derived neuron models of AS. We also provide critical insight into the mechanisms of imprinted expression of *UBE3A* and ASO-mediated unsilencing of paternal *UBE3A*. Our data strongly suggest that displacement of RNA Pol II following ASO cleavage is fundamental to unsilencing paternal *UBE3A.* Guided by our understanding of this mechanism, we have developed a shRNA as a novel therapeutic approach for unsilencing paternal *UBE3A* in human AS neurons. These findings provide strong support for the theory that transcription of *UBE3A-ATS*, rather than the RNA itself, silences paternal *UBE3A*. More broadly, our findings will inform the future design and application of ASOs and other RNA cutters as viable therapeutics.

## Results

### *UBE3A-ATS* targeting ASOs unsilence paternal *UBE3A* in AS neurons

To determine whether ASOs targeting *UBE3A-ATS* activate paternal *UBE3A* in human neurons, we utilized an AS iPSC line (AS del 1-0)^19^ harboring a large deletion of maternal 15q11-q13, which allowed us to assay expression of *UBE3A-ATS* and *UBE3A* solely from the paternal allele. We previously demonstrated that paternal *UBE3A* is silenced in functionally mature cortical neurons derived from this iPSC line^19^. In order to specifically target *UBE3A-ATS*, we selected two ASOs (ATS-ASO1 and ATS-ASO2) which complement *UBE3A-ATS* in a region of *SNHG14* downstream of *SNORD109B* but not overlapping *UBE3A*. We treated AS del 1-0 iPSC-derived neurons with these two ASOs (ATS-ASOs) and observed a dose-dependent reduction of *UBE3A-ATS* RNA levels (Figure 1A). Concomitant with the reduction of *UBE3A-ATS*, we observed a significant increase in *UBE3A* RNA (Figure 1A). Treatment with either of the two ATS-ASOs at a concentration of 10μM resulted in restoration of *UBE3A* RNA levels to approximately 70% that of normal iPSC-derived neurons (Figure 1C). Higher concentrations of ASOs were observed to have cytotoxic effect and precluded analysis of *UBE3A* levels. We further confirmed that ATS-ASO treatment increased UBE3A protein levels in AS-del 1-0 neurons (Figure 1E). We also performed a time-course to determine how long the effects of ATS-ASOs are sustained in AS-del 1-0 neurons (Figure 1F). Reduction of *UBE3A-ATS* was maintained for at least 7 weeks following a single ASO treatment in ATS-ASO-treated neurons compared to neurons treated with a non-targeting ASO (SCRAM). Similarly, *UBE3A* RNA levels remained elevated in ATS-ASO-treated neurons, although the magnitude of increase over SCRAM-treated neurons reached a maximum at 3 weeks and began to decrease thereafter.

**Figure 1.**
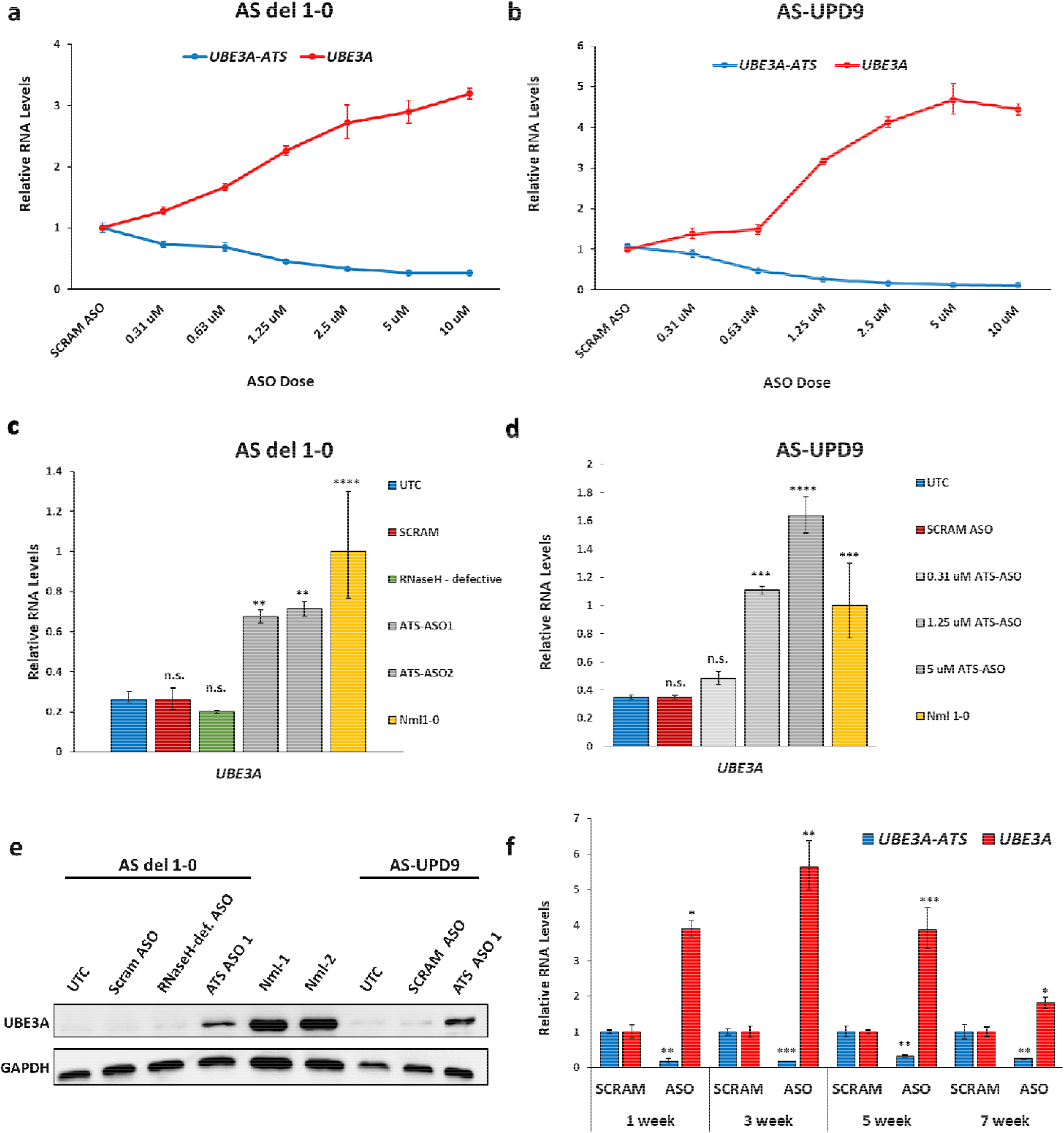
*UBE3A-ATS* targeting ASOs reactivate paternal *UBE3A* in AS iPSC-derived neurons. **a,b** qRT-PCR analysis of *UBE3A-ATS* and *UBE3A* levels in AS del 1-0 and AS-UPD9 iPSC-derived neurons following treatment with increasing doses of ATS-ASO1. RNA levels are presented relative to SCRAM ASO-treated controls. **c, d** qRT-PCR analysis of *UBE3A* levels in AS del 1-0 and AS-UPD9 neurons following treatment with ATS-ASOs or controls (SCRAM, RNaseH-defective). RNA levels are presented relative to neurons derived from a neurotypical control iPSC line (Nml1-0). Error bars depict standard error of the mean of at least three replicates. **p<0.01, ***p<0.001, ****p<0.0001 (Ordinary one-way ANOVA with Dunnett’s multiple comparisons test to UTC) **e** Western blot analysis of UBE3A and GAPDH in AS del 1-0 and AS-UPD9 ASO-treated neurons and neurotypical control iPSC-derived neurons (Nml1 and Nml2). **f** qRT-PCR analysis of *UBE3A-ATS* and *UBE3A* levels in AS del 1-0 iPSC-derived neurons harvested at 1,3,5,and 7 weeks following a single treatment with 10μM ATS-ASO1. RNA levels are presented relative to respective SCRAM ASO-treated neurons at each time point. Error bars depict standard error of the mean of at least three replicates. *p<0.05, **p<0.01, ***p<0.001 (unpaired t-test).

Individuals with AS due to paternal uniparental disomy (UPD) have two silenced paternal *UBE3A* alleles. Unsilencing both paternally-inherited *UBE3A* copies could potentially yield higher than normal levels of *UBE3A*. We generated and characterized iPSCs from an individual with AS due to UPD and verified imprinted *UBE3A* expression upon differentiation into forebrain cortical neurons (Supplemental Figure 1D). Similar to AS del 1-0 neurons, ATS-ASOs reduced *UBE3A-ATS* and increased *UBE3A* in a dose-dependent manner in AS-UPD neurons (Figure 1B). Notably, we found that *UBE3A* RNA levels were restored to near normal levels at much lower ASO doses in AS-UPD neurons. In fact, treatment of AS-UPD neurons with ASO at 5μM and 10μM resulted in approximately 1.6-fold increase in paternal *UBE3A* over normal levels (Figure 1D). UBE3A protein levels in AS-UPD neurons treated with 10μM ATS-ASO were also significantly increased compared to untreated or SCRAM ASO-treated controls (Figure 1E).

Our data show that ATS-ASOs can reduce *UBE3A-ATS* and restore expression of *UBE3A* from both paternal alleles in AS-UPD neurons. This suggests that lower doses of ASOs are sufficient to restore normal levels of *UBE3A* in AS-UPD than in AS due to *UBE3A* deletion. Consideration of the genotype of AS individuals when dosing *UBE3A-ATS*-targeting ASOs is important since an increased level of *UBE3A* may contribute to chromosome 15q duplication syndrome (Dup15q), although it is not clear whether increased *UBE3A* alone is sufficient to disrupt neural development.

### Unsilencing of paternal *UBE3A* rescues physiological phenotypes in AS neurons

We next wanted to test the ability of ATS-ASOs to rescue cellular phenotypes in AS iPSC-derived neurons. AS del 1-0 neurons were treated with ATS- or SCRAM ASOs at 7 weeks of maturation and again at 10 weeks, which provided sustained *UBE3A* beginning at the early stages of differentiation from neural progenitor cell to neuron (Figure 2A). Whole-cell patch clamp recording was performed on neurons at 11 weeks of maturation and the same cultures were subsequently harvested for analysis of *UBE3A* and *UBE3A-ATS* levels by qRT-PCR (Figure 2B). AS neurons were previously shown to have a more depolarized resting membrane potential (RMP) and delayed maturation of action potential (AP) firing patterns compared to neurotypical controls^20^. ATS-ASO-treated neurons displayed a corrected, more hyperpolarized RMP when compared to SCRAM-treated controls (Figure 2C). Analysis of AP firing patterns also showed that *UBE3A* rescue led to significantly more mature APs when compared to SCRAM controls (Figure 2D). Examination of several properties of individual APs across conditions, including amplitude, threshold, and width also supported a restoration of more mature APs in *UBE3A* rescue neurons compared to SCRAM controls (Figure 2E,F,G).

**Figure 2.**
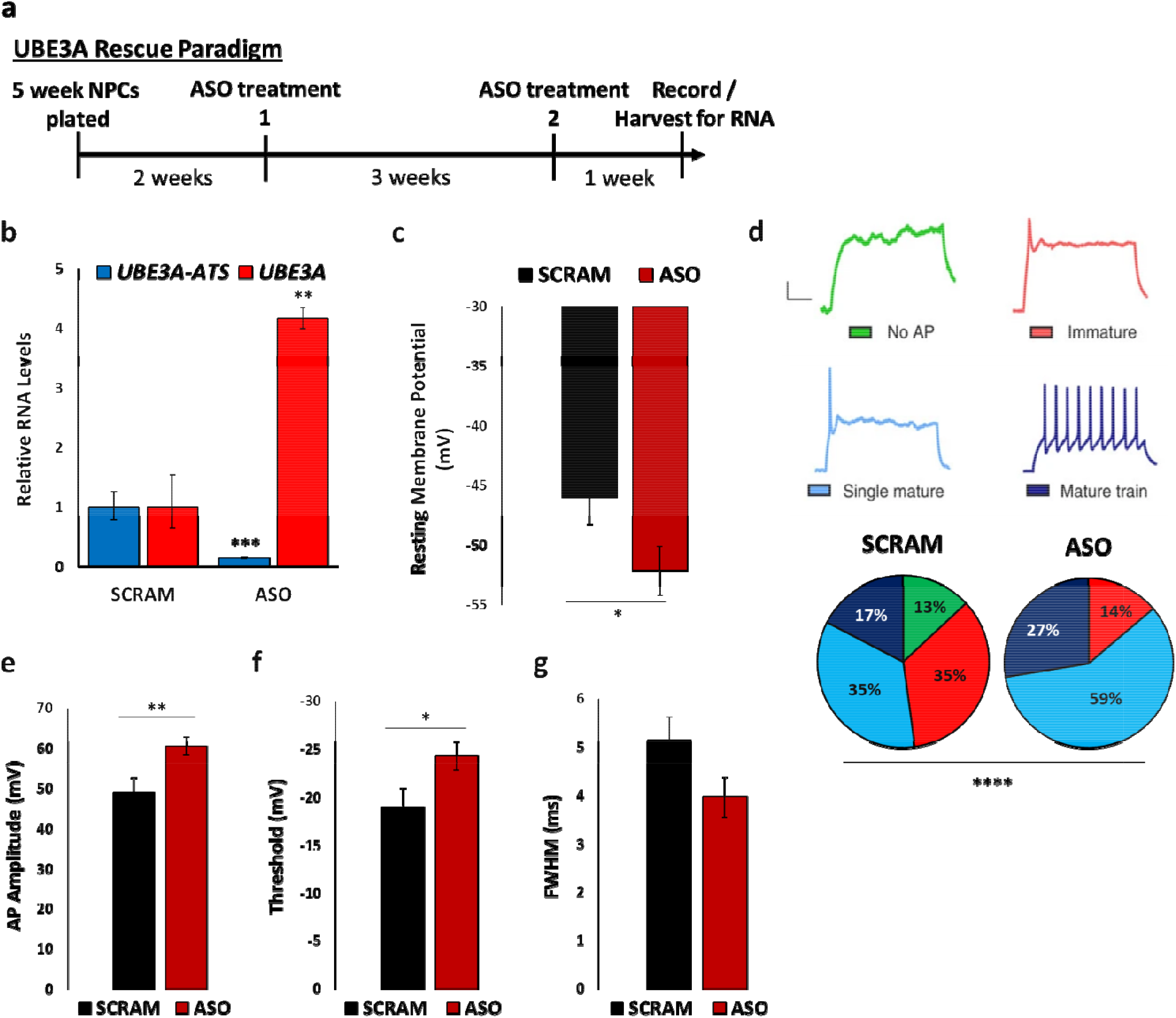
ASO-mediated reactivation of paternal *UBE3A* rescues the disrupted neuronal maturation phenotype in AS iPSC-derived neurons. **a** Timeline of *UBE3A* rescue experiment depicting ASO treatment, patch-clamp recording, and collection of RNA. **b** qRT-PCR analysis of *UBE3A-ATS* and *UBE3A* levels in AS del 1-0 iPSC-derived neurons following treatment with 10μM ATS-ASO1. RNA levels are presented relative to SCRAM-treated controls. Error bars represent standard error of the mean of at least three replicates. **p<0.01, ***p<0.001 (unpaired t-test). **c** Mean resting membrane potential in SCRAM ASO-treated and ATS-ASO-treated AS del 1-0 neurons. Error bars depict standard error of the mean. (n=23 cells for SCRAM and n=29 cells for ATS-ASO-treated neurons in all panels) *p<0.05 (unpaired t-test). **d** (top) Representative action potential (AP) traces in AS del 1-0 iPSC-derived neurons used for categorizing AP patterns in *UBE3A* rescue experiments. (bottom) Distribution of AP patterns in SCRAM controls and ATS-ASO treated neurons. ****p<0.0001 (Chi-square test). **e** Mean AP amplitude **f** threshold, and **g** full-width at half-maximal amplitude (FWHM) in AS del 1-0 neurons following treatment with SCRAM control or ATS-ASO. Error bars depict standard error of the mean. *p<0.05, **p<0.01 (unpaired t-test).

Taken together, these data suggest that restoration of *UBE3A* by ATS-ASOs corrects physiological phenotypes in human AS neurons. The highly specific and dose-dependent ability of ASOs targeting *UBE3A-ATS* to restore *UBE3A* and rescue AS phenotypes in human AS neurons lends strong support for the successful use of ASOs in the treatment of AS.

### ATS-ASOs do not affect the proximal *SNHG14* transcript or processed snoRNAs

*UBE3A-ATS* is located at the distal end of *SNHG14*, a long non-coding RNA that hosts several small nucleolar RNAs (snoRNAs)^21^. These include the *SNORD116* snoRNA cluster, loss of which causes many features of Prader-Willi syndrome^22,23^. Therapeutic approaches to restore paternal *UBE3A* should ideally leave the proximal portions of the *SNHG14* long-non coding RNA unchanged. In order to examine the effect of ATS-ASOs on *SNHG14,* we performed stranded total RNA-seq and qRT-PCR on AS-del 1-0 neurons treated with ATS- and SCRAM control ASOs. We observed no significant reduction in RNA-seq reads mapping to proximal *SNHG14* in the region containing the *SNORD116* snoRNA cluster (Supplemental Figure 2A). qRT-PCR using probe-primer sets recognizing the coding exons of *SNRPN* and the *SNORD116* host gene (*SNORD116HG*) confirmed this (Supplemental Figure 2B). A moderate reduction in reads was detected on the plus strand in the distal portion of *SNHG14*, near *SNORD109B,* in ATS-ASO-treated neurons when compared to SCRAM ASO-treated and untreated AS neurons (Supplemental Figure 2A). Reads mapping to the *SNORD115* cluster are difficult to interpret in these data because the repetitive nature of the sequence prevents accurate mapping of the reads. However, qRT-PCR showed a significant reduction of the *SNORD115* host gene (*SNORD115HG*; Supplemental Figure 2B). Importantly, a significant reduction in reads along the most distal portion of *SNHG14* containing *UBE3A-ATS* and across the region which overlaps *UBE3A* on the plus strand was only observed in neurons treated with the two ATS-ASOs (Supplemental Figure 2A). This reduced expression was also confirmed using qRT-PCR (*UBE3A-ATS*; Supplemental Figure 2B).

The *SNORD116* and *SNORD115* host genes are so-called because they host individual snoRNA copies within their introns. Upon splicing, the functional snoRNAs themselves are folded and bound by ribonucleoproteins before being processed from the introns^24^. Due to the bound proteins, the functional snoRNAs can have long half-lives. We also analyzed the effect of *UBE3A-ATS* ASOs on individual processed snoRNAs by qRT-PCR (Supplemental Figure 2B). We quantified select snoRNAs from the *SNORD116* and *SNORD115* clusters to determine whether the steady-state levels of these more stable RNA species are reduced following ASO treatment. We observed no significant differences in expression of *SNORD116-29*, *SNORD115-1*, *SNORD115-37*, or *SNORD115-48* in ATS-ASO-treated neurons compared to SCRAM ASO-treated controls (Supplemental Figure 2C). Our data shows that ATS-ASOs neither alter expression of the PWS-associated *SNORD116* transcripts nor processed *SNORD116* and *SNORD115* snoRNAs.

### ATS-ASOs terminate transcription of *UBE3A-ATS*

ASOs that unsilence paternal *Ube3a* in mouse and human AS neurons require recruitment of RNase H and cleavage of the ASO:RNA heteroduplex. ASOs modified to be resistant to RNase H recognition did not result in increased *UBE3A* expression^11^ (Figure 1C). Recent work has shown that ASOs which induce RNase H-mediated cleavage of target RNAs can cause transcription termination downstream of the cleavage site via displacement of RNA Pol II^17,18^. Disengagement of RNA Pol II occurs via recruitment of the 5’-3’ exonuclease, XRN2, to the unprotected 5’ end of the RNase H-cleaved RNA, in a process known as the torpedo model for transcriptional termination^25^.

We hypothesized that silencing of paternal *UBE3A* is dependent on the process of transcription of *UBE3A-ATS*, rather than on the RNA itself. Thus, ASOs that unsilence *UBE3A* must disengage RNA Pol II transcribing *UBE3A-ATS*, thereby allowing RNA Pol II transcribing *UBE3A* to complete transcription. To test this hypothesis, we performed precision nuclear run-on and sequencing (PRO-seq) analysis to detect the location of RNA Pol II transcribing *UBE3A-ATS* (plus strand) and *UBE3A* (minus strand) in AS neurons which had been treated with ATS-ASOs or SCRAM controls. The density of RNA Pol II (represented as reads per base) was quantified using a sliding window along the distal portion of *SNHG14* that contains the *UBE3A-ATS*. We observed a striking decrease in the density of RNA Pol II downstream of the ASO cut site (Figure 3A,B). This reduction of RNA Pol II along *UBE3A-ATS* was coupled with a significant increase in RNA Pol II density transcribing *UBE3A* (Supplemental Figure 3). Moreover, ASOs targeting the *SNORD115* cluster caused displacement of RNA Pol II from the plus strand further upstream of *UBE3A-ATS* and a similar increase in RNA Pol II transcribing *UBE3A* (Figure 3A,B; Supplemental Figure 3). The *SNORD115*-ASO also robustly increased paternal *UBE3A* RNA and protein (Supplemental Figure 4A,B).

**Figure 3.**
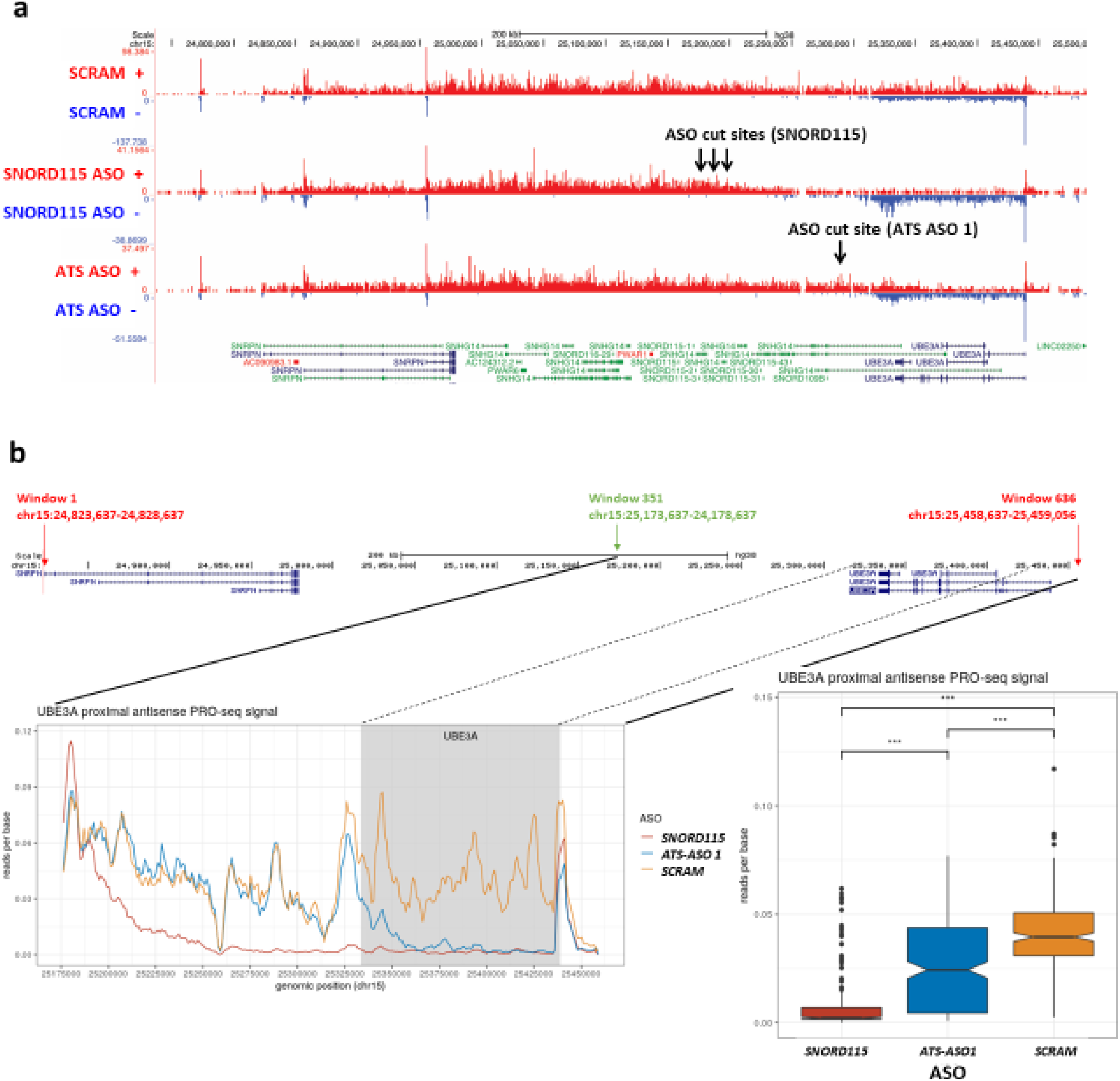
ASO-directed cleavage of *UBE3A-ATS* results in transcription termination and relieves transcriptional interference-mediated silencing of *UBE3A*. **a** Mapped PRO-seq reads on the plus strand (red) and minus strand (blue) across the region of chromosome 15 containing *SNHG14* and *UBE3A* in SCRAM control, SNORD115-, and ATS-ASO-treated AS del 1-0 neurons. Arrows depict ASO target sites. **b** Sliding window analysis of PRO-seq reads along *UBE3A-ATS* in SCRAM control, SNORD115-, and ATS-ASO treated AS del 1-0 neurons. (left) Plot of mapped reads per base across *UBE3A-ATS*. Gray area indicates region where *UBE3A-ATS* overlaps *UBE3A*. (right) Quantification of reads per base across windows 351 through 636 in SCRAM, SNORD115- and ATS-ASO-treated neurons. ***p_SCRAM-SNORD115_<2×10^−16^, p_SCRAM-ATS-ASO1_<4.5×10^−15^, p_SNORD115-ATS-ASO1_<2×10^−16^ (unpaired t-test).

### Transcriptional termination of *UBE3A-ATS* is critical for unsilencing *UBE3A*

Disengaging RNA Pol II not only terminates transcription, but also reduces the steady-state levels of RNA, making it difficult to disentangle the role of the transcribing polymerase versus the *UBE3A-ATS* RNA in repressing paternal *UBE3A*. We sought to find an ASO that would reduce *UBE3A-ATS*, but not terminate transcription. The snoRNA, *SNORD109B,* is located at the distal end of *SNHG14* between the *SNORD115* cluster and *UBE3A*. As discussed above, snoRNAs are bound by ribonucleoproteins (snoRNPs) while the protected snoRNA is processed from introns. Wu et al^26^ showed that the snoRNP-bound snoRNA explicitly protects the 5’ end of RNAs from XRN2-mediated degradation. We hypothesized that *SNORD109B* would prevent XRN2 from terminating transcription via the torpedo model following ASO-mediated RNA cleavage (Figure 4A). To test this hypothesis, we designed an ASO targeting a sequence immediately upstream of *SNORD109B* and assayed *UBE3A* expression. In both AS del 1-0 and AS-UPD9 neurons, although the *SNORD109B*-ASO reduced *UBE3A-ATS* and *SNORD115*, we observed a significantly dampened activation of *UBE3A* compared to the ATS-ASOs (Figure 4B,C).

**Figure 4.**
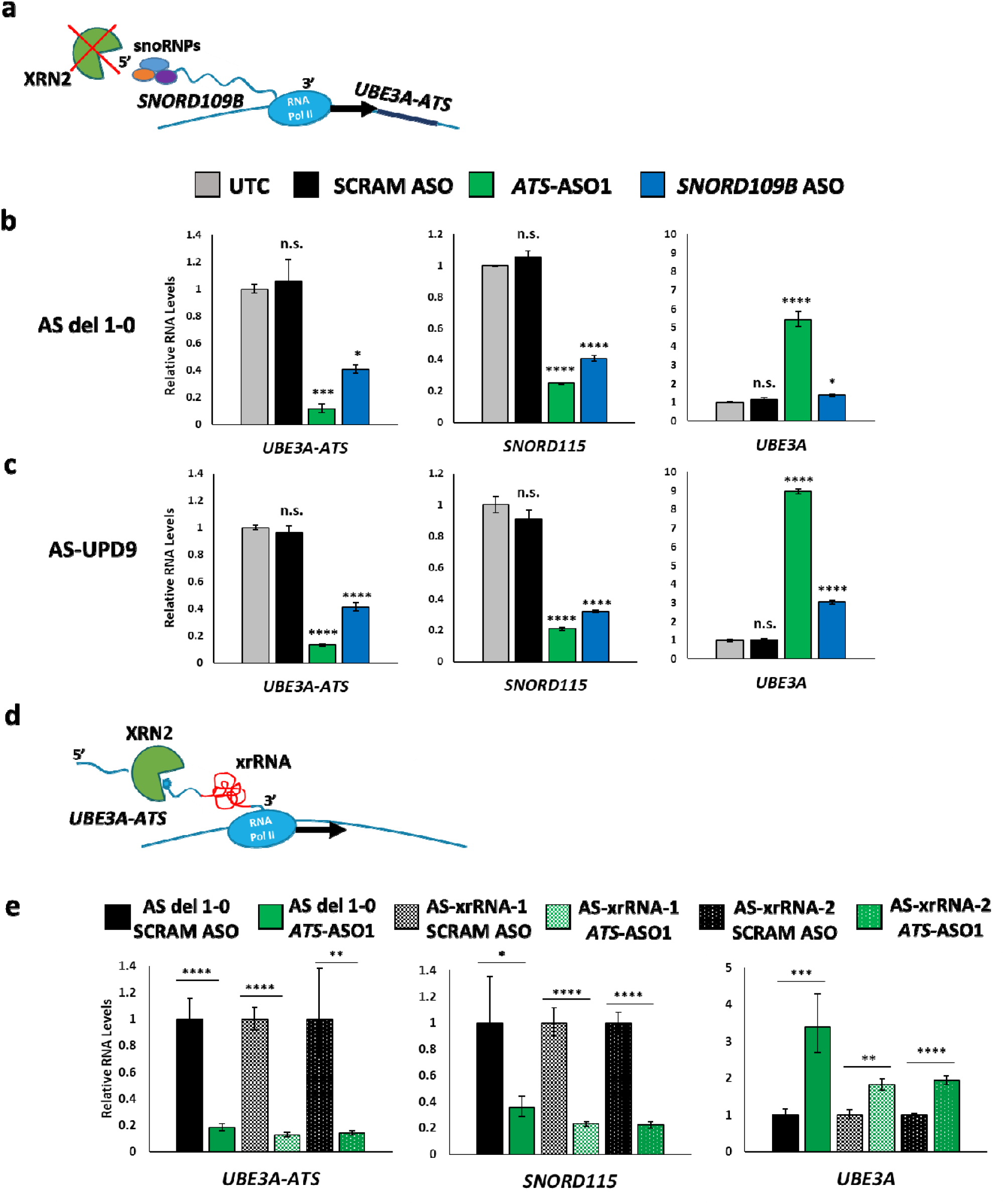
XRN2 displacement of RNA Pol II following ASO-directed cleavage is fundamental to restoration of *UBE3A* expression. **a** Schematic illustrating snoRNP-mediated protection of *SNORD109B* from degradation by XRN2 and prevention of RNA PolII displacement following ASO-directed cleavage. **b, c** qRT-PCR analysis of *UBE3A-ATS*, *SNORD115*, and *UBE3A* levels in AS del 1-0 and AS-UPD9 neurons following treatment with SCRAM, ATS-, and SNORD109B-ASOs. RNA levels are presented relative to untreated controls (UTC). All error bars depict standard error of the mean of at least three replicates. *p<0.05, ***p<0.001, ****p<0.0001 (Ordinary one-way ANOVA with Dunnett’s multiple comparisons test to UTC). **d** Schematic illustrating blockage of XRN2-mediated displacement of RNA PolII following ASO-directed cleavage by insertion of an XRN-resistant RNA pseudoknot (xrRNA) into *UBE3A-ATS*. **e** qRT-PCR analysis of *UBE3A-ATS*, *SNORD115*, and *UBE3A* levels in AS del 1-0, AS-xrRNA-1, and AS-xrRNA-2 neurons following treatment with SCRAM or ATS-ASO. RNA levels are presented relative to SCRAM control ASO-treated neurons for each cell line. All error bars depict standard error of the mean of at least three replicates. *p<0.05, **p<0.01, ***p<0.001, ****p<0.0001 (unpaired t-test).

We also used CRISPR to insert a XRN2-resistant pseudoknot RNA (xrRNA) within *UBE3A-ATS* in AS del 1-0 iPSCs (Supplemental Figure 5). This pseudoknot was previously identified in the zika virus genome, and was shown to be resistant to degradation by XRN1 and XRN2^27^. We hypothesized that the pseudoknot would block XRN2-mediated RNA degradation and therefore prevent disengagement of RNA Pol II following ATS-ASO treatment (Figure 4D). We generated two independent cell lines (AS-xrRNA-1 and AS-xrRNA-2), each harboring the xrRNA sequence in a different location within *UBE3A-ATS* (Supplemental Figure 5A). Upon neural differentiation, these AS-xrRNA neurons were treated with ATS-ASOs or a SCRAM control. AS-xrRNA neurons treated with the ATS-ASO showed a strong reduction of *SNORD115* and *UBE3A-ATS*, as previously observed, but showed a dampened activation of *UBE3A*, much like the *SNORD109B* ASO-treated AS neurons (Figure 4E).

These data support the hypothesis that XRN2 displacement of RNA Pol II following ASO-directed cleavage is fundamental to restoring *UBE3A* expression. They also demonstrate that RNA binding proteins and/or sequence variants may protect RNAs from XRN2-mediated degradation and thus render ASOs less effective at terminating transcription.

### Cleavage of *UBE3A-ATS* by shRNA unsilences paternal *UBE3A*

Based on our understanding of the mechanism by which ASOs targeting *UBE3A-ATS* unsilence paternal UBE3A, we hypothesized that other RNA cleavers may function similarly. To test this, we designed shRNAs to direct cleavage of *UBE3A-ATS* adjacent to the target sites of the ASOs discussed above. Indeed, treatment of AS del 1-0 neurons with *UBE3A-ATS* shRNA significantly reduced *UBE3A-ATS* levels and restored expression of *UBE3A* when compared to a SCRAM non-targeting shRNA (Figure 5A).

**Figure 5.**
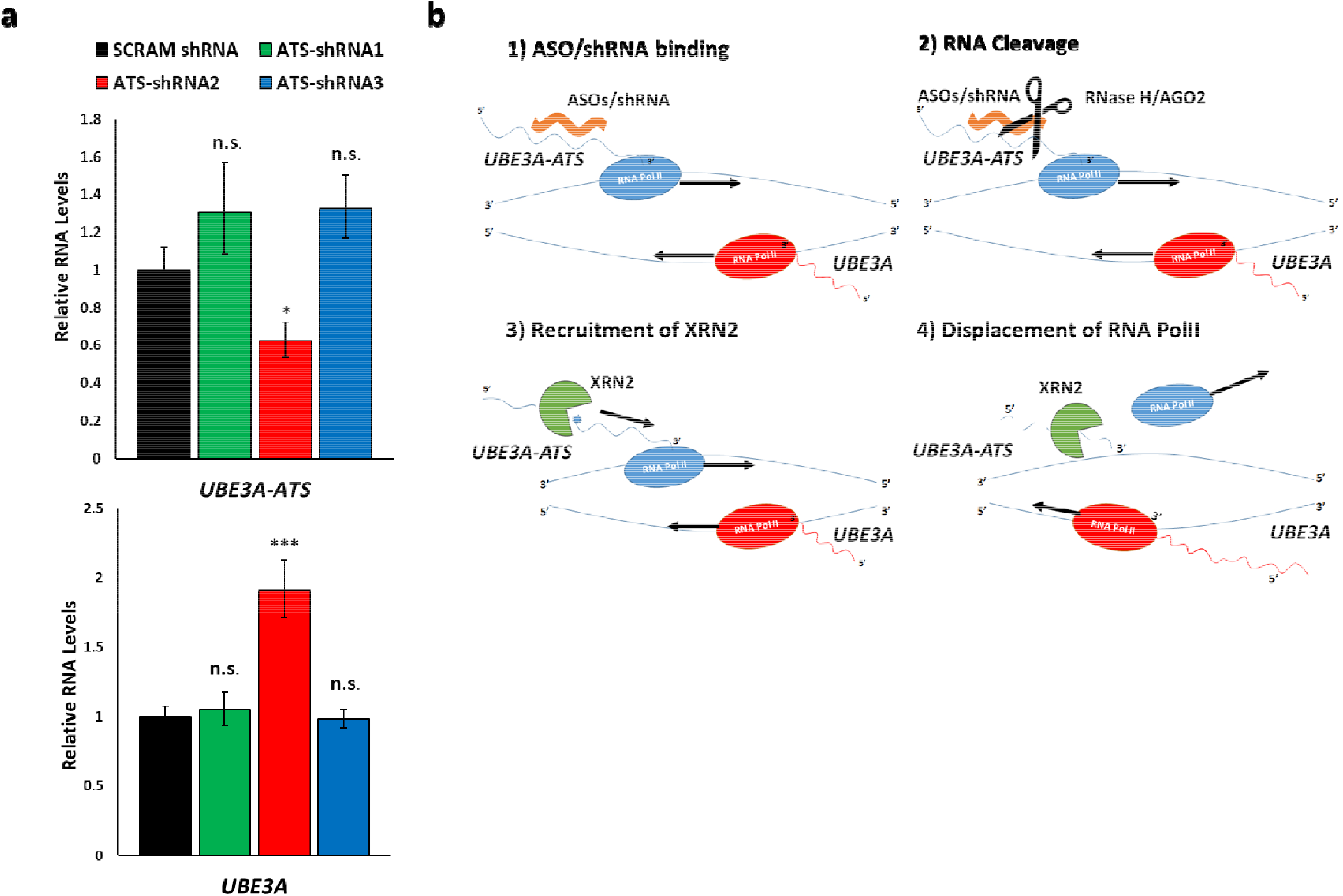
Cleavage of *UBE3A-ATS* by shRNA unsilences paternal *UBE3A.* **a** qRT-PCR analysis of *UBE3A-ATS* (top) and *UBE3A* (bottom) levels in AS del 1-0 neurons following treatment with SCRAM and ATS-shRNAs. RNA levels are presented relative to SCRAM shRNA controls. Error bars depict standard error of the mean of at least three replicates. *p<0.05, ***p<0.001 (unpaired t-test). **b** Proposed model for the mechanism of ASO- and shRNA-mediated unsilencing of *UBE3A*. (1) ASOs/shRNAs bind to *UBE3A-ATS* RNA as it is actively being transcribed. (2) The ASO-*UBE3A-ATS* RNA heteroduplex is cleaved by RNase H. shRNAs cleave UBE3A-ATS via AGO2 (3) The newly cleaved, uncapped *UBE3A-ATS* RNA is degraded by XRN2, which ultimately “catches” and displaces RNA Pol II transcribing *UBE3A-ATS* downstream of the ASO cut site. (4) Removal of RNA Pol II from *UBE3A-ATS* then allows RNA Pol II to complete transcription of *UBE3A*.

We propose the following model for a mechanism of RNA cleavage-mediated unsilencing of *UBE3A* in human neurons (Figure 5B). ASOs and shRNAs first cleave *UBE3A-ATS*. The newly cleaved, uncapped *UBE3A-ATS* RNA is degraded in the 5’ to 3’ direction by XRN2, which eventually “catches” and displaces RNA Pol II downstream of ASO binding on the plus strand. This, in turn, relieves transcriptional interference and allows RNA Pol II to continue transcribing *UBE3A* from the minus strand, restoring functional *UBE3A* levels.

Our mechanistic understanding of *UBE3A* imprinting and its ASO-mediated reversal provides crucial insights into other potential AS therapeutics. shRNAs, like those described here, can be delivered using viral vectors, such as AAV, enabling a single-dose treatment for restoring *UBE3A* expression in AS. Other RNA cutters, including Cas13 and trans-cleaving ribozymes may provide similar opportunities for therapeutic development, potentially offering improved therapeutic options for Angelman syndrome.

## Conclusion

We show here that ASOs direct RNase H-mediated cleavage of *UBE3A-ATS* and unsilence *UBE3A* in human AS neurons. Restored *UBE3A* in these AS neurons is sufficient to rescue electrophysiological phenotypes. We demonstrate that ASO-mediated cleavage of *UBE3A-ATS* results in displacement of RNA Pol II transcribing *UBE3A-ATS* several kilobases downstream of ASO binding, strongly suggesting that ASOs terminate transcription of *UBE3A-ATS via* the torpedo model. Furthermore, failure to terminate transcription of *UBE3A-ATS* was associated with a dampened activation of *UBE3A*. These data provide the most comprehensive support for the transcriptional interference hypothesis for silencing of paternal *UBE3A* and provide critical mechanistic insight into the function of ASO-based therapeutics for Angelman syndrome.

## Materials and Methods

### iPSC derivation and culture

AS-UPD9 iPSCs were derived from donated patient peripheral blood by the University of Connecticut Cell and Genome Editing Core. The patient blood sample was obtained following informed consent under UConn Health IRB protocol number 15-174-1 and was deidentified prior to reprogramming. We have complied with all required ethical regulations. Episomal vectors containing the Yamanaka reprogramming factors were used for reprogramming according to published protocols. Loss of heterozygosity along the entire long arm of chromosome 15 in AS-UPD9 iPSCs was confirmed by Illumina CytoSNP-850K v1.2 array by the University of Connecticut Chromosome Core. AS del 1-0 iPSCs were generated as previously described^19^. AS-xrRNA-1 and AS-xrRNA-2 iPSCs were engineered by using CRISPR/Cas9 (as previously described^9,28^) to insert the sequence of an Xrn1-resistant RNA pseudoknot from Zika virus downstream of the location of ATS-ASO binding in AS del 1-0 iPSCs. Small guide RNAs were designed to two locations within the distal end of SNHG14 to direct Cas9 cutting. A DNA repair template was provided in the form of a single-stranded oligonucleotide (ssODN) which contained the RNA pseudoknot sequence flanked by regions of homology to the sequence adjacent to the CRISPR cut sites. Positively targeted iPSC clones were identified by PCR of genomic DNA using primers flanking the CRISPR cut site and subsequently confirmed by Sanger sequencing. CRISPR sgRNA, ssODN, and primer sequences are provided in Supplemental Table 2.

iPSCs were maintained on irradiated mouse embryonic fibroblasts in human embryonic stem cell medium which consists of DMEM/F12, 20% knockout serum replacement, 1mM L-glutamine, 1X nonessential amino acids, 100mM β-mercaptoethanol (all Gibco products through Life Technologies), and 8ng/mL basic fibroblast growth factor (bFGF, Millipore). iPSCs were manually passaged by cutting and pasting colonies every 6 or 7 days.

### Methylation Analysis

Analysis of methylation at the Prader-Willi syndrome imprinting center (PWS-IC) was performed as previously described.^19^

### Neural differentiation

iPSC-derived neural progenitors were generated by monolayer differentiation according to established protocols^29^. After five weeks of neural differentiation neural progenitors were plated on poly-D-lysine/laminin-coated substrates in neural differentiation medium (NDM) consisting of Neurobasal Medium, B-27 supplement, nonessential amino acids, and L-glutamine (all Gibco products through Life Technologies) supplemented with 1μM ascorbic acid, 200μM cAMP, 10ng/mL brain-derived neurotrophic factor (BDNF, Peprotech), and 10ng/mL glial-derived neurotrophic factor (GDNF, Peprotech) and allowed to mature in culture for several weeks. All experiments were conducted on neural cultures that were at least 10 weeks old.

### ASO and shRNA design and treatment

ASOs targeting *UBE3A-ATS*, *SNORD115*, and a non-targeting control (SCRAM) were provided by Ionis Pharmaceuticals. The ASO targeting adjacent to *SNORD109B* was designed and purchased through Integrated DNA Technologies. ASOs were synthesized as previously described^11^ and were 20 bp in length, with five 2’-O-methoxyethyl-modified nucleotides at each end of the oligonucleotide, ten DNA nucleotides in the center and a phosphorothioate backbone. ASOs were added to cultured neurons in antibiotic-free NDM at a final concentration of 10μM except as noted in dose-response curve experiments. shRNAs were designed using The Broad Institute Genetic Perturbation Platform tools (https://portals.broadinstitute.org/gpp/public/) and cloned into the pLKO.1-puro vector. Lentiviral particles were produced from cloned shRNAs in HEK293 cells and used to transduce cultured neurons. ASO sequences are provided in Supplemental Table 1.

### qRT-PCR

Total RNA was isolated from iPSCs or iPSC-derived neurons using RNA-Bee (AMS Biotechnology) according to the manufacturer’s protocol. cDNA was produced using the High Capacity cDNA Reverse Transcription Kit (Life Technologies).

Analysis of 15q11-q13 genes in iPSCs and iPSC-derived neurons was performed at least in triplicate. All qPCR assays used were TaqMan Gene Expression Assays (Life Technologies). Ct values for each gene were normalized to the house keeping gene *GAPDH*. Relative expression was quantified as 2^^−ΔΔCt^ relative to the calibrator sample. Data are presented as the mean relative expression plus or minus the standard error of the mean (of ΔCt).

### Western blotting

Neurons were processed for protein analysis as previously described^20^. The following antibodies were used: rabbit anti-UBE3A (Bethyl laboratories, 1:1000), mouse anti-GAPDH (Millipore, 1:2000), and goat anti-mouse and goat anti-rabbit HRP-conjugated secondary antibodies (Cell Signaling Technologies, 1:3000).

### PRO-seq library preparation

PRO-seq libraries were generated using 2 × 10^6^ permeabilized cells as described in Mahat et al.^30^ and Judd et al.^31^, with the following modifications. 25,000 permeabilized Drosophila S2 nuclei were added as a spike-in control during the run-on reaction. After the run-on reaction, RNA was extracted using Norgen RNA purification columns (Cat #37500) as per the manufacturer’s protocol. Following purification, RNA was base hydrolyzed for 20 min on ice prior to enrichment of biotinylated nascent RNA using DynaBeads MyOne Streptavidin C1 (Invitrogen, Cat #65001). 3’ RNA adapter ligation was carried out off-bead using 25 pmol adapter, followed by 5’ decapping, 5’ hydroxyl repair, and 5’ RNA adapter ligation performed on beads. Upon completion of reverse transcription, libraries were pre-amplified for a total of 5 cycles using cycling parameters from Mahat et al^30^. Test amplifications using serial dilutions of the pre-amplified libraries were then performed to determine the ideal number of cycles for full-scale amplification, with 12 cycles chosen for all samples. Fully amplified libraries were purified using NEB Monarch PCR & DNA cleanup kits (Cat # T1030L) or by PAGE purification on an 8% PAGE gel (ATS-ASO sample only). Final libraries were quantified by Qubit, pooled in an equimolar fashion, and submitted for sequencing on an Illumina NextSeq 500 at the Center for Genome Innovation (UConn, Storrs, CT).

### PRO-seq data mapping

Adapter sequences (*TGGAATTCTCGGGTGCCAAGG*) were trimmed from raw reads using *cutadapt*^32^, filtered for a minimum length of 15bp, and trimmed to a maximum of 36bp. Reads were then reverse complimented using FASTX-Toolkit^33^. Reads were first aligned to the human rDNA repeat (GenBank: U13369.1) using Bowtie^34^ with the -k 1 and -v 2 options. Unaligned reads were then mapped to a concatenated human (GRCh38/hg38) and Drosophila (dm6) genome with the -v 2 and -m 1 options to filter for unique alignments. Aligned reads were converted to the bam format using SAMtools^35^, then bed files of the 3’ terminal base of each read were generated using Bedtools^36^ in order to represent the position of RNA polymerase II. These 3’-end bed files were used for all downstream analysis. Lastly, bedgraphs used for data visualization were generated by calculating the coverage of the 3’ position of reads and normalized to the read depth of the Drosophila spike-in controls.

### Sliding window analysis of PRO-seq data

Sliding window analysis was used to assess transcription levels across the *UBE3A* locus in response to ASO treatments. For analysis in the sense orientation, a custom annotation for *UBE3A* was generated to include a region 5kb upstream and 20kb downstream of the longest annotated isoform from GRC27. Sliding windows 5kb long with 1kb overlaps were generated using *bedtools makewindows* with the -reverse option as *UBE3A* is on the minus strand. For analysis in the antisense orientation, a region spanning from the TSS of the upstream, antisense SNRPN gene to 20kb past the 5’ end of *UBE3A* was used. As with the sense orientation, 5kb sliding windows with a 1kb overlap were generated using bedtools, though without the -reverse option. Counts in sense and antisense windows that overlapped the proper strand were calculated using *bedtools coverage*. Counts data were imported into R 3.5.0^37^, normalized to reads per base, then used to calculate fold change relative to the scrambled control ASO. Data were tidied using the *tidyr* and *dplyr* packages^38^ then plotted using the *ggplot2^38^*, *ggpubr*, *ggsci*, and *ggsignif* packages. Line plots depict the center of each window.

### Electrophysiology

Whole-cell voltage and current clamp recordings of 10- to 13-week-old stem cell derived neurons were performed as previously described^20^. Briefly, recordings were performed on individual coverslips in a recording chamber fixed to the stage of the microscope that was continuously perfused with oxygenated artificial cerebrospinal fluid (aCSF). Neurons were identified based on their morphology when viewed on infrared differential interference contrast (DIC) video microscopy. Patch pipettes (3 to 5 MΩ) were pulled from borosilicate glass capillaries and filled with internal solution containing KCl, K-gluconate, HEPES, phosphocreatine, EGTA, CaCl^2^, Na^2^-ATP and Na-GTP. Input resistance (Ri) was noted throughout the recording period. Resting membrane potential (RMP) was noted at break-in by injection of 0 current. RMP values were corrected for liquid junction potential offline (JPcalc). To elicit AP firing, cells were held in current clamp mode at approximately −70 mV, followed by application of 500 ms current steps from −20 to +40 pA in intervals of 5 pA. Offline analysis of data was done using Clampfit software.

### Total RNA Sequencing

Total RNA was isolated from ASO-treated and untreated control AS del 1-0 neurons using RNA-Bee (AMS Biotechnology) according to the manufacturer’s protocol. Ribosomal RNA-depleted stranded RNA libraries were prepared using the TruSeq Stranded Total RNA Prep Kit (Illumina) and 500ng of total RNA as input. Libraries were multiplexed and sequenced on a NextSeq 500 sequencer (Illumina) to generate 75bp paired-end reads. Reads were aligned to the human genome (hg19) using Bowtie (version 2.1.0.0) and Tophat (version 2.0.8)^34^. At least 20 million mapped reads were generated for each sample.

## Acknowledgements

We thank the members of the Chamberlain, Core, and Levine labs for discussion of the data presented here and review of this manuscript. We also acknowledge the University of Connecticut Cell and Genome Engineering Core for generation of iPSC lines used in this study.

## Author Contributions

NDG and SJC conceived of and designed the study, collected and analyzed data, and wrote the manuscript. DG performed and analyzed electrophysiology experiments and generated the AS-xrRNA iPSC lines. RD and LC performed and analyzed PRO-seq experiments. PJ, AW, and FR designed and provided ASOs. ESL assisted with electrophysiology analysis and provided critical feedback during preparation of the manuscript.

## Data and materials availability

RNA-seq data are available through the NCBI Sequence Read Archive (SRA) with accession number XXXXXX. PRO-seq data are available at The Gene Expression Omnibus (GEO) with accession number XXXXXX. Cell lines are available upon reasonable request and after completion of Material Transfer Agreements with the authors through the University of Connecticut Cell and Genome Engineering Core.

