## Supplemental Information for "Antisense oligonucleotides targeting *UBE3A-ATS* restore expression of *UBE3A* by relieving transcriptional interference"

**Supplemental Table 1. ASO sequences.**

|  | **Sequence 5’ 🡪 3’** |
| --- | --- |
| **ASOs** |  |
| SCRAM ASO | CCTTCCCTGAAGGTTCCTCC |
| ATS-ASO 1 | AGTAAGGTCTGTTATTCTCC |
| ATS-ASO 2 | ATCATGTGCATACCCAGGGT |
| *SNORD115* ASO | GTCATCACCTCTCTTCAGGA |
| *SNORD109B* ASO | GTTTCCACAGCCAGGCTTAT |

**Supplemental Table 2. CRISPR guide RNAs, repair template oligos, and screening primers for generation of AS-xrRNA iPSCs.**

|  | **DNA Sequence 5’ 🡪 3’** |
| --- | --- |
| **xrRNA target site 1 guide RNA** | GATCTCATGTTGCCACGGGAG **TGG** |
| **xrRNA target site 2 guide RNA** | GGTATTCGATATACGTATCC **CGG** |
| **xrRNA target site 1 repair template ssODN**  (Zika virus xrRNA sequence highlighted in red) | CCATTGAAGAAAAACCCGCGTTCAAAATGGTAGAAAATTGTATGGCATTTTCACTTGCACTTGCCCTGCCACTCCCGCAGGCTGCACAGCTTTCCCCAAACTGTGGCTGACTAGCAGGCCTGACATGGCAACATGAGATCTTGGTTTGTAGAAGGTGGGAAACCGGTTCCCAATTTTCTCCCTTCAACCA |
| **xrRNA target site 2 repair template ssODN**  (Zika virus xrRNA sequence highlighted in red) | CATGTTAAAGGCCTCCTCAGATTCCTATACTCTATTCTTGCAAAGCTGGAGAGGTTCTTTAGAAGTATTTTAGCCGGGACAGGCTGCACAGCTTTCCCCAAACTGTGGCTGACTAGCAGGCCTGACATACGTATATCGAATACCCTCAACTTCACCAATGCAAGCCAGCTTCTAACATACTAATTAGCAAGCCACTGATT |
| **xrRNA target site 1 screening primer FWD** | CTGCCACCTTTCCCTATCTG |
| **xrRNA target site 1 screening primer REV** | TGAATCAGCAAAATGGTGGA |
| **xrRNA target site 2 screening primer FWD** | TTGTCATTTCACAGGTCTCCA |
| **xrRNA target site 2 screening primer REV** | CAGCAATAGGCTCTGCTGTG |

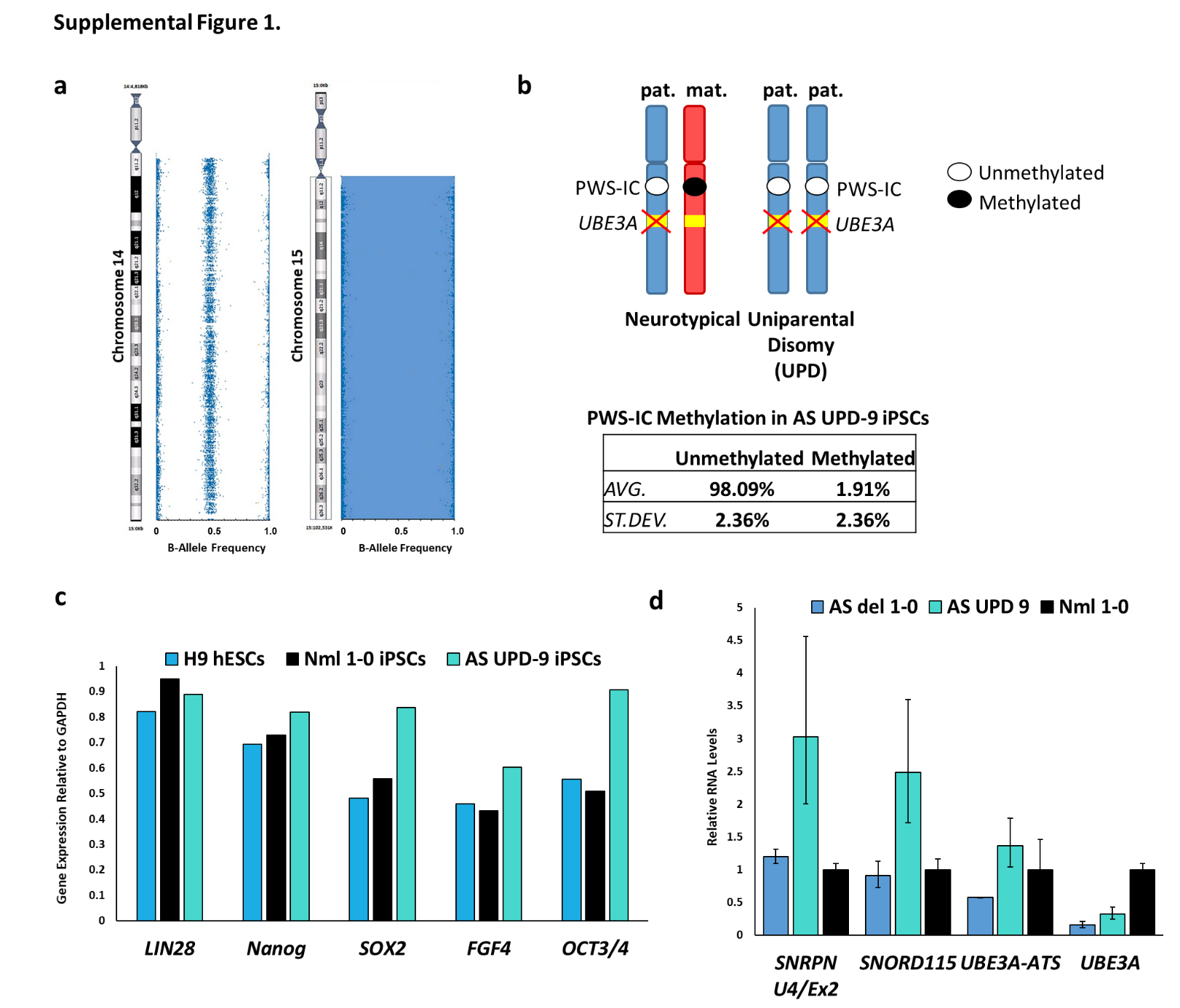

**Supplemental Figure 1. Characterization of AS paternal uniparental disomy (AS-UPD) iPSCs. a** B-Allele Frequency graphs for chromosomes 14 and 15 show loss of heterozygosity along the entire q arm of chromosome 15 in AS-UPD9 iPSCs. b. (top) Schematic depicting the methylation status at the Prader-Willi syndrome imprinting center (PWS-IC) on maternal (mat.) and paternal (pat.) chromosome 15 in neurotypical and AS-UPD individuals. Unmethylated DNA at the PWS-IC is represented by an open circle. Methylated DNA is represented by a closed circle. Table shows results of methylation analysis in AS-UPD9 iPSCs. Methylation-dependent restriction digest and qPCR was performed in duplicate on separate genomic DNA samples. **c** qRT-PCR analysis of selected pluripotency genes in AS UPD-9 iPSCs. H9 hESCs and Nml1-0 iPSCs are included for reference. **d** qRT-PCR analysis of *SNRPN U4/Ex2*, *SNORD115HG*, *UBE3A-ATS*, and *UBE3A* in AS del 1-0, AS-UPD9, and Nml1-0 iPSC-derived neurons. RNA levels are presented relative to Nml1-0 neurons. Error bars depict standard error of the mean of at least three replicates.

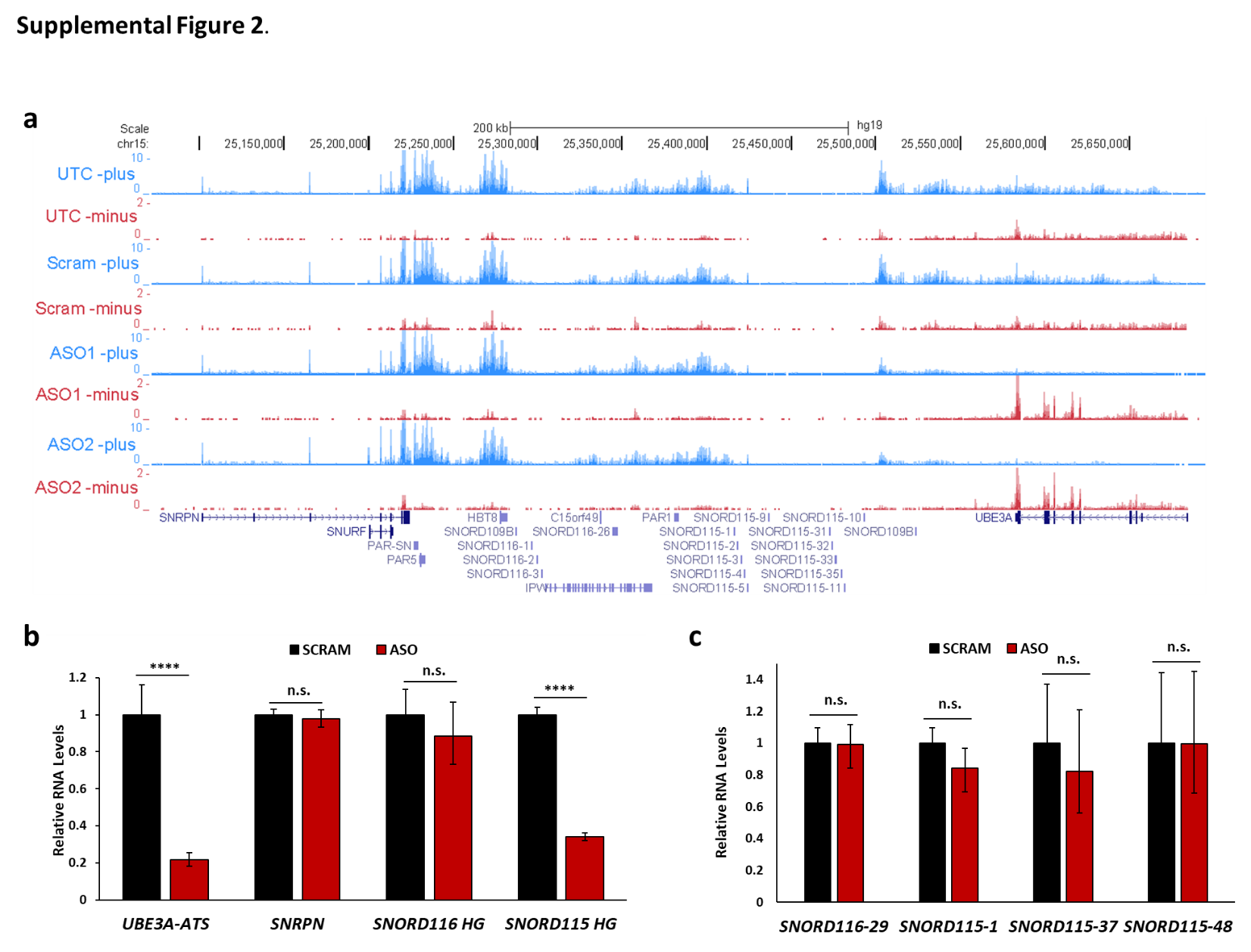

**Supplemental Figure 2**. **ASOs targeting *UBE3A-ATS* do not affect the proximal *SNHG14* transcript or its processed snoRNAs. a** Total RNA-seq reads mapped to the plus strand (blue) and minus strand (red) in the region of chromosome 15 containing *SNHG14* and *UBE3A* in untreated control (UTC), SCRAM-, ATS-ASO1-, and ATS-ASO2-treated AS del 1-0 neurons. **b** qRT-PCR analysis of *UBE3A-ATS*, *SNRPN*, *SNORD116* host gene (*SNORD116HG*) and *SNORD115* host gene (*SNORD115HG*) in SCRAM control and ATS-ASO treated AS del 1-0 neurons. RNA levels are presented relative to SCRAM ASO-treated neurons. ****p<0.0001 (unpaired t-test). **c** qRT-PCR analysis of selected snoRNAs, which are processed from the *SNORD116* and *SNORD115* host gene transcripts, in SCRAM control and ATS-ASO treated AS del 1-0 neurons. RNA levels are presented relative to SCRAM ASO-treated neurons. All error bars depict standard error of the mean of at least three replicates.

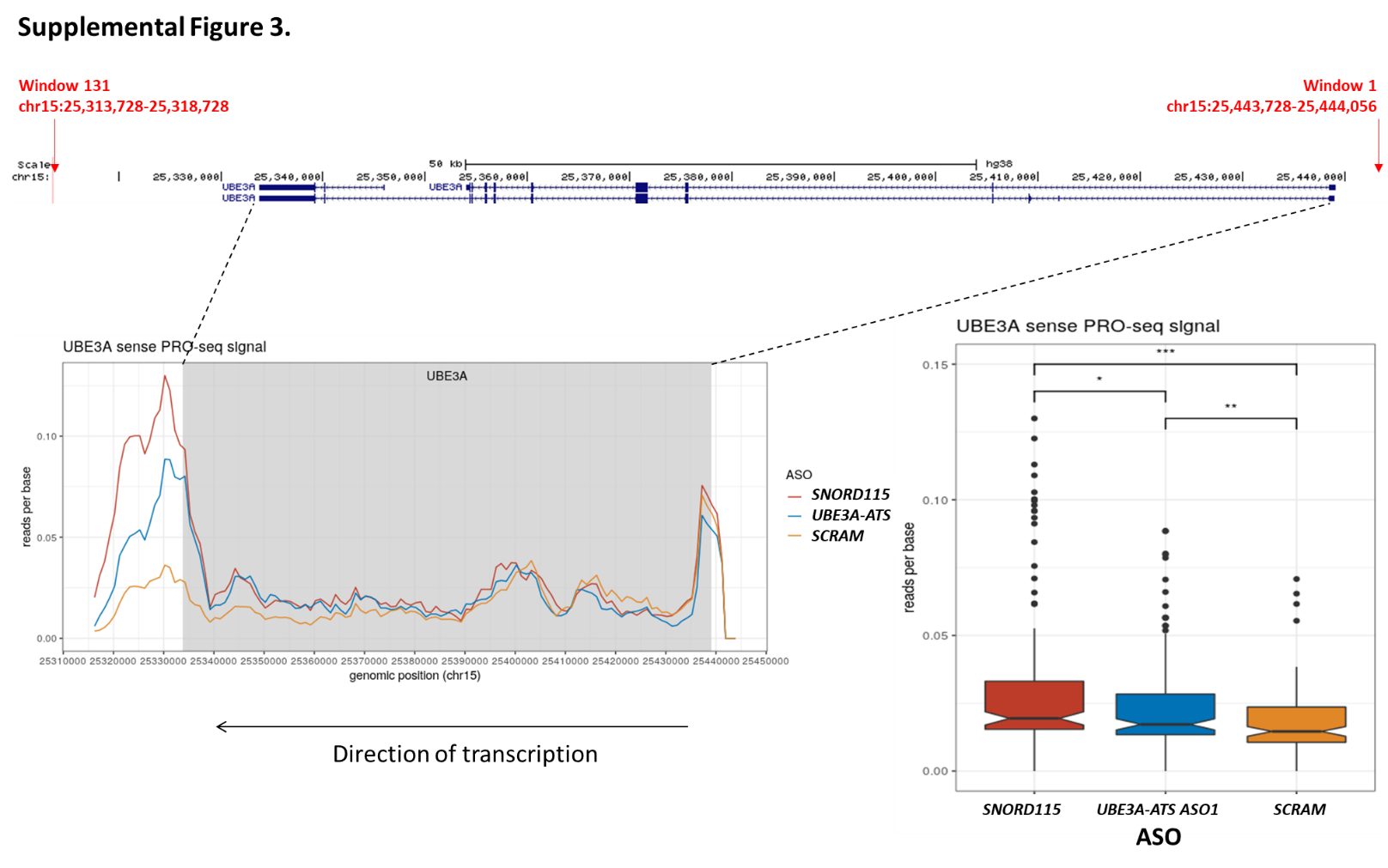

**Supplemental Figure 3. PRO-seq analysis of *UBE3A* in ASO-treated AS del 1-0 neurons.** Sliding window analysis of PRO-seq reads along *UBE3A* in SCRAM control, SNORD115-, and ATS-ASO treated AS del 1-0 neurons. (left) Plot of mapped reads per base across *UBE3A*. Gray area indicates the span of annotated UBE3A gene, as indicated by dotted lines (right). Quantification of reads per base across all 131 windows in SCRAM-, SNORD115- and ATS-ASO-treated neurons. *p<0.05, **p<0.01, ***p=1.6X10^-6^ (unpaired t-test).

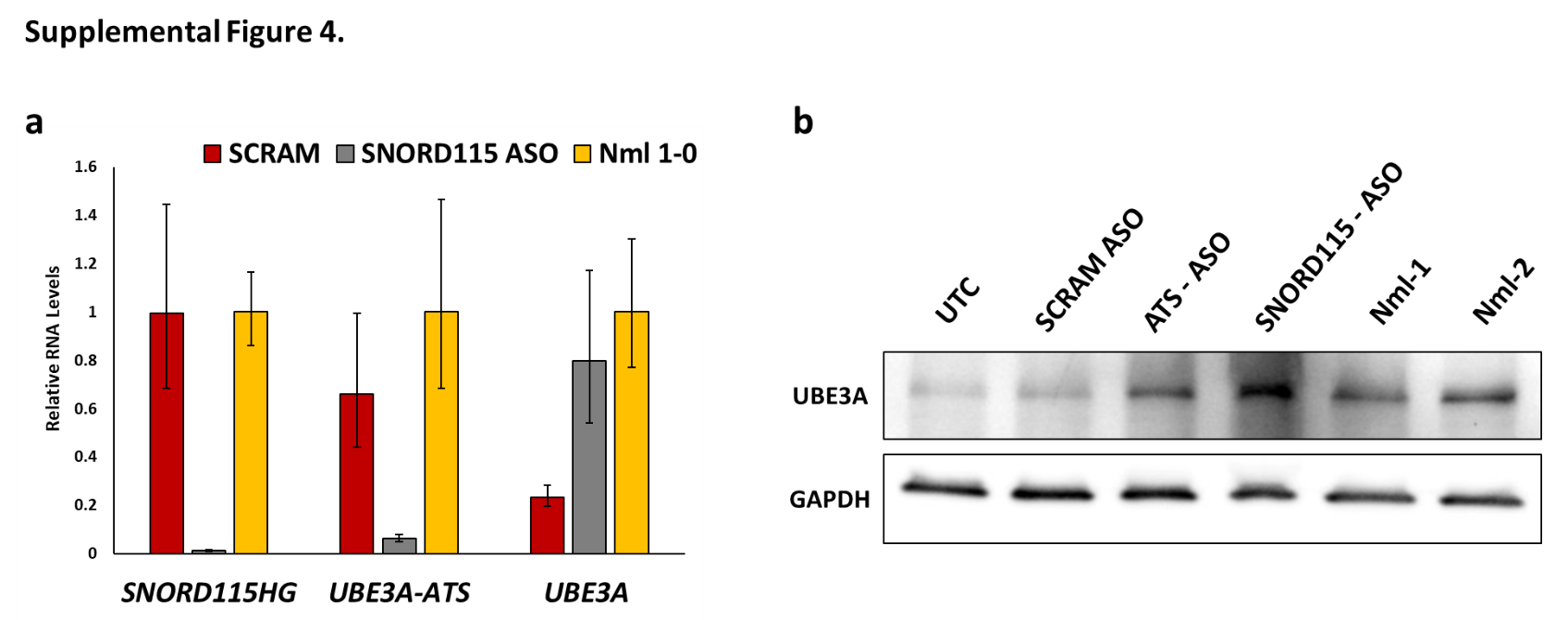

**Supplemental Figure 4. ASOs targeting *SNORD115* reactivate paternal *UBE3A* in AS neurons. a** qRT-PCR analysis of *SNORD115HG*, *UBE3A-ATS*, and *UBE3A* in AS del 1-0 neurons following treatment with SCRAM control or SNORD115-ASOs. RNA levels are presented relative to Nml1-0 control neurons. Error bars depict standard error of the mean of at least three replicates. **b** Western blot analysis of UBE3A and GAPDH in AS del 1-0 neurons treated with SCRAM-, SNORD115-, and ATS-ASOs and neurotypical control neurons (Nml1 and Nml2).

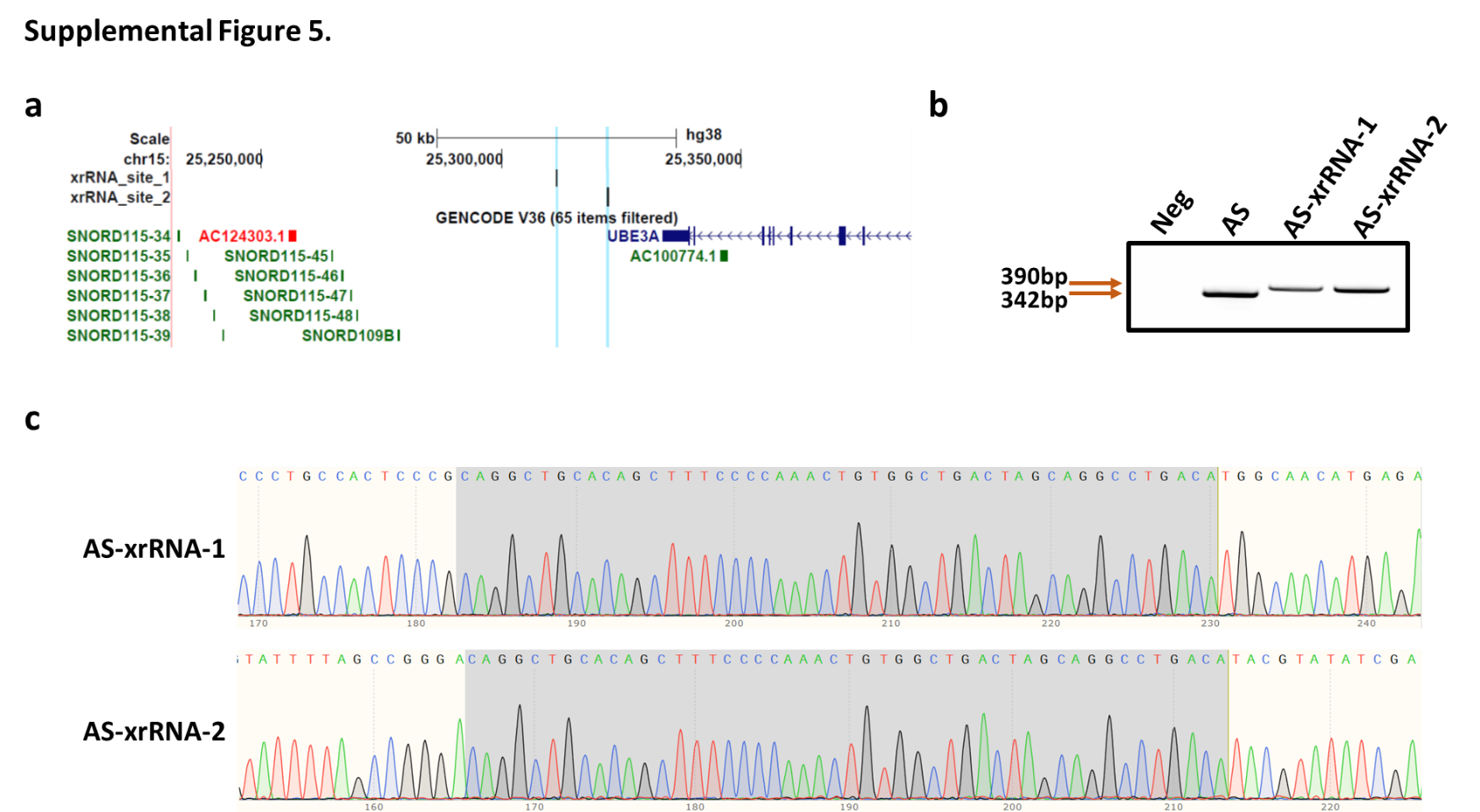

**Supplemental Figure 5. Generation of AS-xrRNA iPSCs. a** Genomic position of the two XRN-resistant RNA (xrRNA) insertion sites (blue bands). **b** PCR of genomic DNA with primers spanning the CRISPR cut site confirm insertion of the 48 base pair (bp) xrRNA in two clones (AS-xrRNA-1 and AS-xrRNA-2). Neg = no DNA control. **b** Sanger sequencing of AS-xrRNA-1 and AS-xrRNA-2 iPSC genomic DNA. Gray area indicates the inserted xrRNA sequence.
